# Females facilitate male patch discovery in a wild fish population

**DOI:** 10.1101/478537

**Authors:** Lysanne Snijders, Ralf H. J. M. Kurvers, Stefan Krause, Alan N. Tump, Indar W. Ramnarine, Jens Krause

## Abstract

When individuals are more socially responsive to one sex than the other, the benefits they get from foraging socially are likely to depend on the sex composition of the social environment. We tested this hypothesis by performing experimental manipulations of guppy, *Poecilia reticulata*, sex compositions in the wild. Males found fewer novel food patches in the absence of females than in mixed-sex compositions, while female patch discovery did not differ between compositions. We argue that these results were driven by sex-dependent mechanisms of social association: Markov chain-based fission-fusion modeling revealed that males reduced sociality when females were absent, while less social individuals found fewer patches. Females were similarly social with or without males. Finally, males, but not females, preferred to join females over males at patches. Our findings reveal the relevance of considering how individual and population-level traits interact in shaping the adaptive value of social living in the wild.

## Introduction

Social living is ubiquitous in nature and has important ecological consequences (Krause & Ruxton 2002; Danchin *et al.* 2004; Guttal & Couzin 2010; Kurvers *et al.* 2014; Gil *et al.* 2018). To thoroughly understand the evolution and maintenance of social living it is important to study the role of social behavior in fitness relevant contexts in the wild, such as foraging. Many animals forage socially (Giraldeau & Caraco 2000), yet individuals seldom respond to all conspecifics equally and there is ample evidence that animals preferentially associate with some conspecifics more than others (Croft *et al.* 2005; Formica *et al.* 2011; Kerth *et al.* 2011; Mourier *et al.* 2012; Kurvers *et al.* 2014; Shizuka *et al.* 2014). Such non-random social associations are expected to influence an individual’s foraging success. Indeed, social associations predicted if and where individuals found new food patches in several species in the wild (Aplin *et al.* 2012; Jones *et al.* 2017; Schakner *et al.* 2017; Snijders *et al.* 2018).

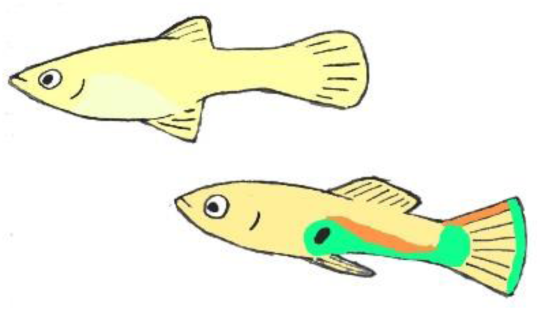

Sex-biased social affinity is one of the most basic and widespread forms of preferential social association in nature and can have important fitness consequences (Silk *et al.* 2003; Cameron *et al.* 2009). Next to social affinity, animals also show preferred social avoidance, for example, by specifically avoiding males to elude male aggression or sexual harassment (Galliard *et al.* 2005; Darden & Croft 2008; Galezo *et al.* 2018). When individuals are indeed more socially attracted and/or responsive to one sex than the other, individual benefits of social foraging are likely to depend on the sex ratio in the local environment. Yet, studies that manipulate the adult sex ratio of foragers, while staying under the selective pressures of the wild, are extremely rare, especially in vertebrate systems. Due to the complexity of effectively manipulating social composition under natural conditions, the few studies that link adult sex ratio to male and female foraging dynamics in the wild are primarily correlational (e.g. Choudhury & Black 1991; Madden *et al.* 2009; Tettamanti & Viblanc 2014, but see Magurran & Seghers 1994). Given that individuals in the wild self-select their social environment (e.g. Croft *et al.* 2003b), drawing any causal inferences without experimental evidence is problematic, which hampers a thorough understanding of the factors that determine the adaptive value of social foraging and, consequently, social living.

Here, we performed a unique manipulation of the social environment using wild individually-marked guppies, *Poecilia reticulata*. This species lives in a fission-fusion society with adult sex ratios fluctuating heavily in time (i.e. season) and space (Pettersson *et al.* 2004). During the dry season, wild guppies naturally form ephemeral subpopulations in temporarily isolated pools (Magurran 2005), what allowed us to manipulate subpopulation adult sex ratios in a natural context without the risk of individuals re-distributing themselves. We formed single-sex (male and female) and mixed (50:50) sex compositions (18 in total) and introduced these to natural pools in the rainforest in Trinidad. In these pools, we studied the individuals’ social behavior, which we analyzed with an individual-based dynamic modeling approach using first-order Markov chain (MC) models (Wilson *et al.* 2014; Krause *et al.* 2017; Snijders *et al.* 2018). Subsequently, we examined individual success in locating novel (experimentally introduced) food patches.

With this study we provide a direct experimental test of how two basic biological factors, individual sex and sex composition, interact to affect individual foraging success in the wild. First, we tested whether sex composition affected an individual male’s and female’s patch discovery success. Second, we tested whether sex composition affected the time an individual male or female spent social and, third, we analyzed if this time spent social directly correlated to patch discovery success. Finally, we examined if there was evidence for sex-biased social attraction before and during the foraging trials. Given that females are typically the preferred association partner for both sexes, we predicted males to find more and females to find fewer food patches in mixed compared to single-sex compositions and that these patterns could be explained by both males and females spending more of their time socially when (more) females are present.

## Material & Methods

### Study area

Our study took place in the Upper Turure rainforest region (10°41’8”N, 61°10’22”W) of Trinidad (11-25 March 2017). The study area was located upstream and received little sunlight, likely making guppies relatively food-limited (Grether *et al.* 2001). Furthermore, the area has few guppy predators present (i.e. ‘low-predation’). We conducted our study in three natural pools (approximate surface area range of the pools: 3-6 m^2^). The in- and outflow of these pools was slightly altered to prevent fish migration but a continuous flow was maintained. All guppies originally occurring in the pools were removed.

### Study subjects

Fish from nearby pools were used as experimental subjects. After capture, fish were sexed, sized and individually marked using an established method of elastomer coloring (VIE marks) (Croft *et al.* 2003a, 2004). We used batches of eight fish, either all-male, all-female or mixed-sex (4 males and 4 females). Fish from each batch were caught from the same area, so fish were likely to be familiar with each other. Each sex composition was replicated twice in each pool, resulting in six batches for each of the three sex compositions. We balanced the sex composition (treatment) order across pools, with pool 1 starting with all-male, pool 2 with all-female, and pool 3 with mixed sex. In total, we thus used 18 groups of eight fish (*N* = 144). After marking, fish were released in the designated study pool and left overnight to settle prior to the social phenotype observations and the subsequent day of foraging trials. Because a few fish left the observation pool before or after the social phenotype observations, we ended up with slightly varying sample sizes for the social phenotyping and foraging trials (*N* social = 141, *N* foraging = 140, *N* combined = 139; see *Supplementary Tables S1a-b* for details). The fish densities in our experiments were typical for our study area and within the range of Trinidadian guppy population densities (Reznick & Endler 1982).

We performed all research in accordance with the law and animal ethical standards of the country in which the study was performed, Trinidad and Tobago. Specifically, our study protocol adhered to the ‘Basic Principles Governing the Use of Live Animals and Endangered Species in Research at the University of the West Indies’ as part of the ‘Policy and Procedures on Research Ethics’ of the University Committee on Research Ethics. All fish were released at the end of their trials.

### Social phenotypes

To quantify the social dynamics, we performed focal follow observations the day before, and thus independent of, the foraging trials. Between 09:00 and 15:00, each fish was followed for two min recording its nearest and second nearest neighbor every 10 sec. A fish was considered a neighbor if it was within four body lengths of the focal fish (Wilson *et al.* 2014; Krause *et al.* 2017). After following each fish for two min, we waited for 5 min upon which we repeated this procedure five more times, resulting in a total of 12 min of focal follows per individual fish in each batch.

### Foraging trials

The day following the social phenotype scoring, we carried out foraging trials during which we presented novel food sources to the guppies. As food source, we used small led balls (8 mm diameter) covered in a mix of gelatin and fish food (TetraPRO©), including carotenoids, a valuable resource for guppies (Kodric-Brown 1989). Within each pool, we introduced the food sources in five locations. Food introduction was standardized by entering the food through small plastic cylinders floating at the water surface (but anchored to the bottom of the pool). The cylinders were open at the top and bottom. The five feedings locations within each pool were roughly evenly distributed over the pool. Upon the start of a trial, we gently lowered the food source in the pool through the plastic cylinder using a monofilament fishing line attached to a wooden rod. Once in the water the food source was kept approximately two centimeters above the ground. Once a fish discovered the food source (i.e. food patch), we waited for one more minute after which we removed the food source and ended the trial. If the food was not discovered within three minutes the trial was also ended. After finishing a trial, we waited for five minutes before starting a new trial. We presented food sources at each of the five locations in a randomized order. Once we completed all five locations, we repeated this procedure four more times, resulting in 25 food presentations per batch (and 450 food presentations in total). After we finished the food presentations for a given batch, all fish of that batch were caught and released further downstream.

### Video analyses

All trials were recorded with Sony Handycams (SONY HDR-PJ530E), mounted on tripods. A single observer used the open-source event-logging software BORIS (Friard & Gamba 2016) (v 4.0) to score for each fish its presence (continuous) during one minute after the arrival of the first fish. An arrival (and subsequent presence) was defined as a fish being within two body lengths of the food source. These data were reviewed by a second observer and cross-referenced with data on discovery time and fish identity collected in the field. If mismatches occurred, trials were double checked and if necessary corrected by a third observer, who had also been present in the field (L.S.). In 423 of the 450 trials (94%) at least one fish found the food resource. However, because of recording problems (e.g. water surface glare), only a subset of the videos was of sufficient quality for reliable identification of all the individuals present (*N* = 391).

### Statistical analyses

#### Generalized linear mixed models

We analyzed the data using generalized linear mixed models (GLMM) with R (R Core Team 2017) version 3.5.1 in R Studio version 1.0.453 (© 2009-2018 RStudio, Inc.), using the *glmer* function in the ‘*lme4’* package (Bates *et al.* 2015) with “bobyqa” as optimizer. Control variables such as body length (as fixed covariate, scaled and centered on sex), individual identity nested in batch identity (as random factor) and pool identity (as fixed factor with three levels) were kept in the model at all times. Individual identity was used as an observation-level random effect in proportional-binomial models to effectively manage overdispersion. Binary-binomial models additionally included trial number (fixed covariate, scaled) and location nested in pool (random factor). To test the significance of fixed effects, we compared models with and without the fixed effect of interest, using Log Likelihood Ratio (LLR) tests. Post-hoc pairwise comparisons (averaged over pools) were made using the R-package ‘*emmeans*’ (Lenth 2018). P-values were adjusted following the Tukey method for comparing a family of four estimates. We based conclusions for individual social traits on permutation models (see below).

#### Markov chain social analyses

To quantify the social dynamics and their potential differences between sex compositions, we used the Markov chain-based fission-fusion model introduced by Wilson et al. (2014). The social behavior of each individual fish is described as a sequence of behavioral (social) states, being either in the presence of a specific conspecific (within four body lengths) or alone. When an individual is with a specific conspecific, it can thus transition to being alone, but it can also stay with this specific ‘nearest’ neighbor or switch to another nearest neighbor [for more details see the Supplementary material of Wilson et al. (2014)]. The data collected during the social phenotype observations were used to estimate the transition probabilities between each state for each individual fish. The overall social propensity (i.e. social time) of a particular fish is subsequently quantified as *P*_a→s_/ (*P*_s→a_ + *P*_a→s_), where *P*_a→s_ is the probability of ending being alone and *P*_s→a_ is the probability of ending a social contact. To examine whether individuals exhibited a sex-bias in their time spent social, we adapted the original model to distinguish between spending time social with male and female nearest neighbors (see Supplementary methods for details). Additionally, we calculated the ϒ-measure to quantify the degree in which individuals express preferences for particular social partners. The ϒ-measure is the sum of squares of the normalized association strengths (relative number of contact moments) between one individual and all others (Boccaletti *et al.* 2006; Krause *et al.* 2017). It was previously shown that guppies occupy consistent positions in social networks regarding both measures (Wilson *et al.* 2015; Krause *et al.* 2017).

#### Model details per research question

##### Does sex composition influence discovery of food patches?

To test our main prediction—that males find more and females find less food patches in mixed-sex batches compared to single-sex batches—we tested whether the interaction between sex and sex composition had a significant effect on individual patch discovery. We quantified individual patch discovery (dependent variable) as binary (yes/no; only including videos of sufficient quality for reliably identifying each individual as present or absent (*N* = 391)). The binary-binomial model included the interaction between sex (male or female) and sex composition (mixed-sex or single-sex) as well as the above-mentioned control variables.

We constructed a proportional-binomial model to test if males and females in mixed-sex batches differed in their likelihood of arriving first at a food patch. As dependent variable, we quantified the number of patches an individual reached being first, divided by the total number of patches reached by its batch. The model included sex as fixed factor and the above-mentioned control factors.

##### Does sex composition influence social time?

To test if males and females differed in the time they spend social depending on sex composition, we used two approaches: a group comparison approach between compositions and a randomization approach within composition (for mixed-sex batches). To test differences in time spent social between males and females within mixed compositions, we permuted (10,000 repetitions) the (highly dependent) social times among males and females within each batch. As test statistic, we used the absolute value of the difference of the mean social times between males and females. However, for the comparison of social times of male-only and female-only groups it is problematic to randomize, because it would involve permutations across groups, mixing up the values of dependent and independent variables. Therefore, we computed the overall social times per batch, which are independent, and used a Wilcoxon rank sum test to compare the values for the female single-sex compositions with those for the male single-sex compositions. The same procedure was applied to the comparison of males(/females) in mixed-sex compositions and males(/females) in single-sex compositions. Furthermore, we analyzed the relationship between social time and body length, by permuting the individual social times within each batch and using Pearson’s correlation coefficient between social time and body length as test statistic (10,000 repetitions).

##### Does social time predict patch discovery?

To examine if variation in individual patch discovery between the four combinations of sex (male, female) and sex composition (single sex, mixed sex) could be explained by corresponding variation in social time, we replaced the factors sex and sex composition with the covariate social time in the binary-binomial model for patch discovery. Significance of social time was evaluated via a randomization procedure (see Supplementary methods for details). We repeated these randomizations 10,000 times and each time calculated the coefficient for social time. We then compared the original model coefficient for social time to the distribution of the coefficients of the permutated models (Farine & Whitehead 2015). Social time and body size were significantly positively correlated in all sex-compositions, i.e. larger fish were more social (Female-only: *coefficient* = 0.48; *P* = 0.001, *N* = 47; Male-only: *coefficient* = 0.24; *P* = 0.02, *N* = 48; Mixed-sex: *coefficient* = 0.33; *P* = 0.03, *N* = 47), yet, we found no evidence of collinearity issues (all Generalized Variance Inflation Factors < 2) and therefore left both social time and body size in the models for patch discovery. Conclusions did not change (effects only became stronger) when we omitted body size or replaced it by the residuals of body size over social time.

##### Is there evidence for sex-specific attraction?

To test if males and females exhibited a sex preference in their social time, we calculated for each mixed-sex batch the percentages of time spent social (based on our Markov model), distinguishing between sex of the focal individual and sex of the social partner. These observed percentages were then compared to percentages based on 10,000 permutations, swapping the identities of social partners. Furthermore, to test for sex-specific attraction during the foraging trials, we tested for mixed-sex compositions whether the sex of the first fish at a food patch predicted the proportion of batch members that would subsequently join. The proportional-binomial model included, for each of the detected trials, the number of fish present divided by the number of subjects in the batch as dependent variable and included sex of the first fish and the before-mentioned control variables as fixed effects. To investigate potential differences in responses between female and male ‘followers’, we ran the same model including only potential (i.e. available) male followers or only potential female followers.

## Results

### Males reach fewer novel patches in absence of females

At a group-level, the male-, female- and mixed-sex compositions were very similar in food patch discovery success, i.e. the proportion of patches found by at least one individual (*Total%* (*Min – Max*)): all-male (*N* = 6): 93% (84 - 100), all-female (*N* = 6): 94% (72 – 100), mixed (*N* = 6): 95% (88-100). On an individual-level, however, fish differed in how likely they were to find a patch, depending on the combination of their own sex and the sex-composition of their social environment (sex*sex composition: *Estimate (Est) ± SE* = −0.80 ± 0.24, *χ^2^* = 10.95, *N* = 3,246, *N* individuals = 140, *P* = 0.0009). Males in mixed compositions were more likely than males in single-sex compositions to reach a new food patch (*Odds ratio ± SE = 1.72 ± 0.29, z-ratio = 3.24, P = 0.007*, Fig. 1), but females in mixed compositions did not differ from females in single-sex compositions (*Odds ratio ± SE = 0.78 ± 0.13, z-ratio = −1.52, P = 0.42*, Fig. 1). Males and females within mixed compositions were equally likely to reach a new food patch (*Odds ratio ± SE = 0.76 ± 0.14, z-ratio = 1.46, P = 0.46*, Fig. 1), while males and females in single-sex compositions differed in the favor of females (*Odds ratio ± SE = 1.68 ± 0.23, z-ratio = 3.75, P = 0.001*, Fig. 1). Individuals were more likely to reach a new food patch over time (i.e. trial number: *Est ± SE*= 0.15 ± 0.04, *χ*^2^= 16.21, *N* = 3246, *N* individuals = 140, *P* < 0.0001). Body size (centered on sex) did not predict the likelihood of patch arrival (*Est ± SE*= 0.10 ± 0.06, *χ*^2^ = 2.83, *N* = 3,246, *N* individuals = 140, *P* = 0.09).

**Figure 1.**
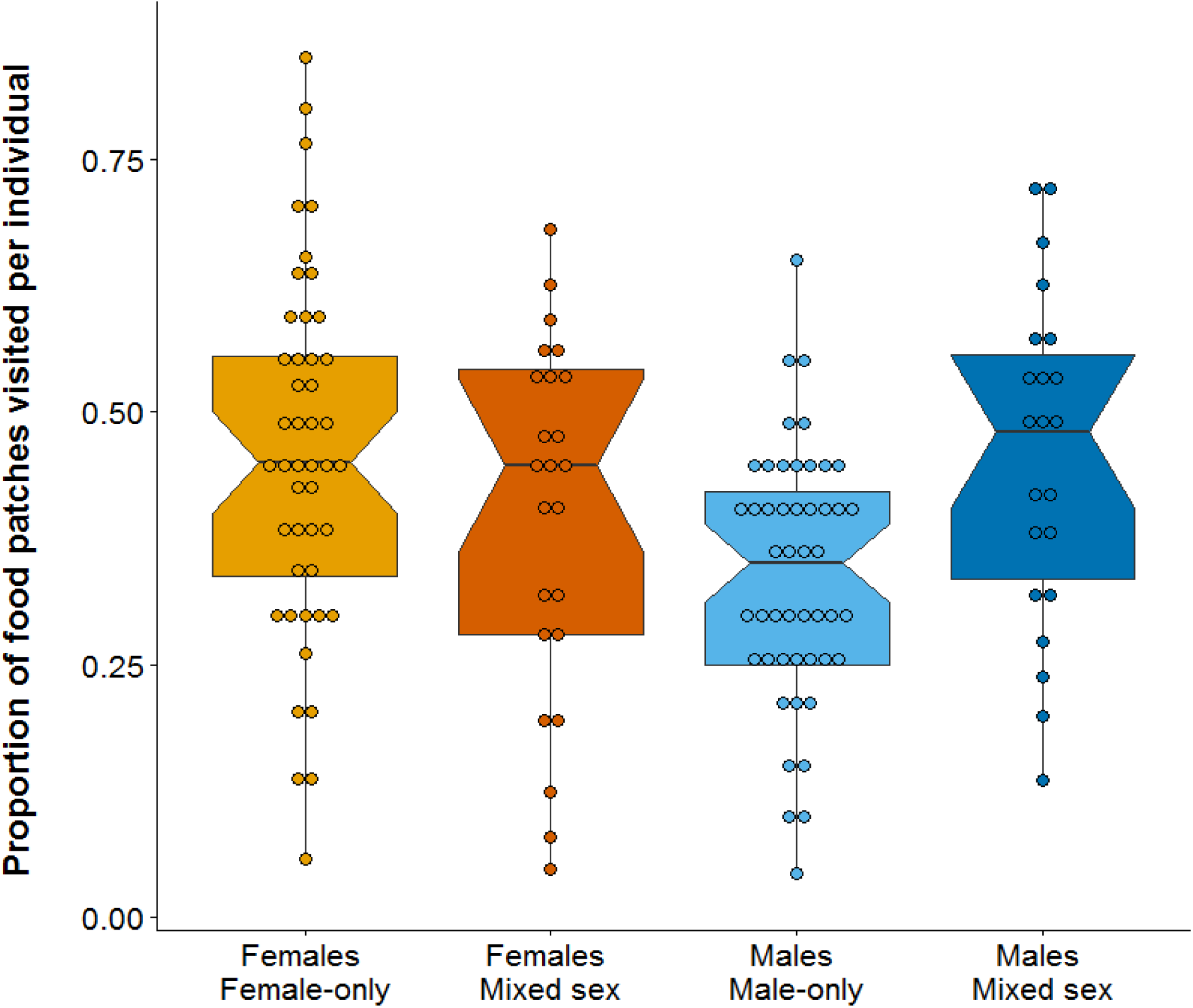
The proportion of novel food patches discovered by individual males and females in relation to sex composition. Males in single-sex compositions reached less food patches than males in mixed-sex compositions, but females in single-sex compositions did not differ from females in mixed-sex compositions. Females in single-sex compositions reached more patches than males in single-sex compositions, yet males and females within mixed-sex compositions did not differ from each other. Box plots show median and 25th to 75th percentiles with whiskers of 1.5 interquartile distances. Non-overlapping notches suggest a significant difference in medians.

Importantly, males and females in mixed compositions did not differ in the proportion of detected patches they were the first to arrive at (*Est ± SE* = −0.16 ± 0.24, *χ^2^* = 0.43, *N* = 46, *P* = 0.51), suggesting that males and females did not differ in their skills to initially detect a food patch (i.e. without social information).

### Males are less social in absence of females

The differences between sex and sex compositions in patch discovery (Fig. 1) closely mirrored differences in individual social time, i.e. the propensity of individuals to spend time near conspecifics (Fig. 2). Males in single-sex compositions spent substantially less time social (14%) than males in mixed-sex compositions (26%, *W* = 34, *P* = 0.009, *N* = 12 groups, Fig 2) and females in single-sex compositions (27%, *Wilcoxon rank sum test*, *W* = 35, *P* = 0.004, *N* = 12 groups, Fig 2). The overall social times of individual females in single-sex compositions and in mixed-sex compositions did not differ (28%, W = 19, P = 0.94, N = 12 groups, Fig 2). Within mixed compositions there was no difference between males and females in the time they spent social (10,000 randomization steps, *score* = 0.022; *P* = 0.47, *N* = 47 individuals, Fig. 2) nor in how they distributed their social contact moment over social partners, i.e. the ϒ-measure (10,000 randomization steps, *score* = 0.004; *P* = 0.62, *N* = 47 individuals). When comparing the underlying social dynamics (i.e., transition probabilities) between the different sex and sex compositions, we found that males in single-sex compositions had a higher likelihood of leaving their nearest neighbor, a higher probability of transitioning from a social state to being alone and a lower probability of transitioning from being alone to being social, compared to individuals in the other sex compositions (Fig. 3, Supplementary Table S2).

**Figure 2.**
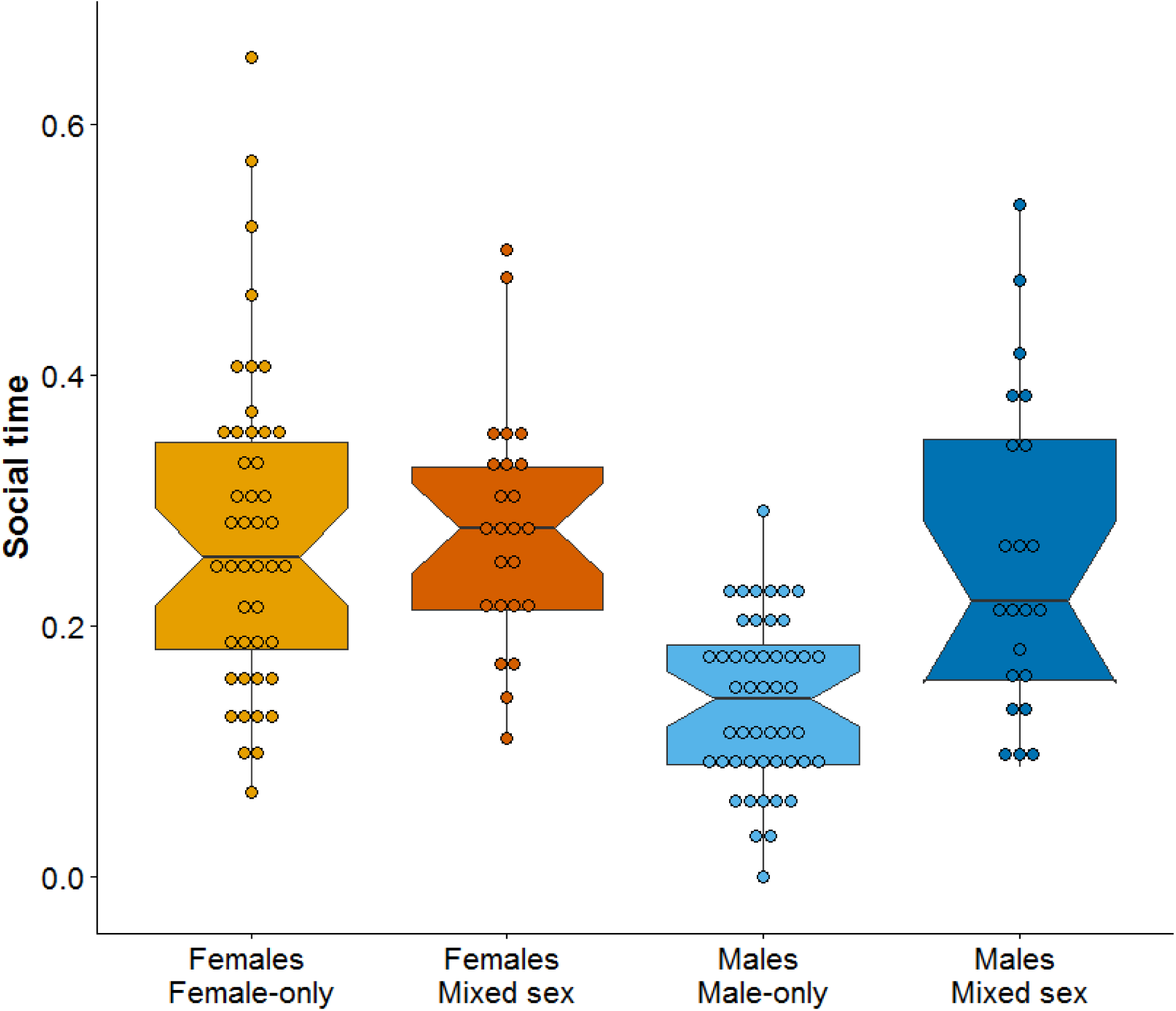
Time spent social by individual males and females in relation to sex composition. A higher Social time value indicates a stronger propensity to spend time in proximity of conspecifics (before the foraging trials). Males in single-sex compositions spent less time social than males in mixed-sex compositions or females in in single-sex compositions. Within mixed-sex treatments, males and females did not differ. Box plots show median and 25th to 75th percentiles with whiskers of 1.5 interquartile distances. Non-overlapping notches suggest a significant difference in medians.

**Figure 3.**
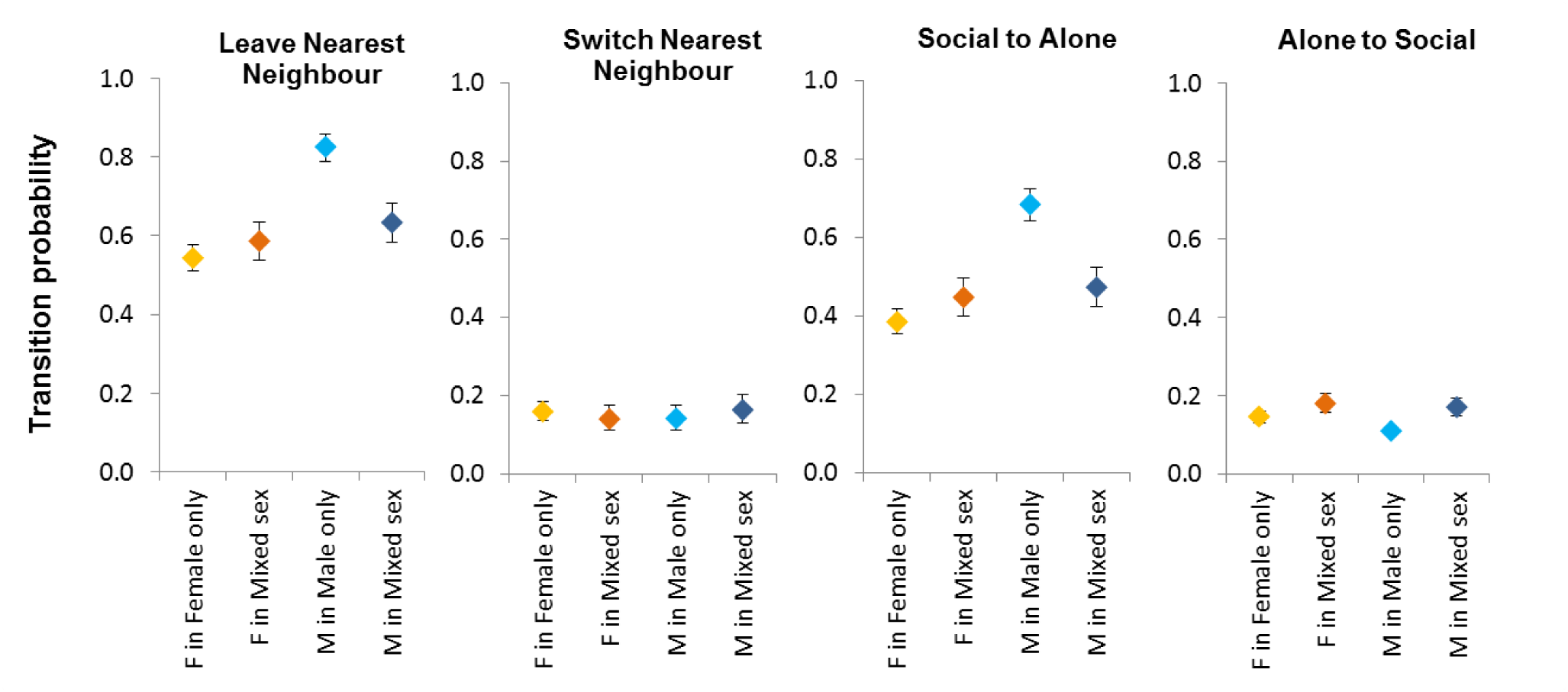
Markov Chain transition probabilities between various social states and being alone in relation to sex composition. Males in male-only compositions (light blue) were more likely to leave their nearest neighbor, more likely to switch from a social state to being alone and less likely to transition from being alone to a social state, than fish in other compositions. There were no differences between sex compositions in the transition probability to switch nearest neighbor. Error bars represent 95% confidence intervals.

### More social individuals reach more novel patches

Differences in individual social time between sex compositions are highly relevant because social time positively predicted the proportion of discovered novel food patches by individuals, across sex compositions (10,000 randomization steps, *coefficient* = 0.22, *N* =3,344, *N* individuals = 144, *P* = 0.0009, Fig. 4). Within sex compositions (Supplementary Table S3, Supplementary Fig. S1), more social males as well as females in mixed-sex compositions reached more patches, while time spent social by males and females in single-sex compositions did not lead to increased patch-discovery. There was also an overall tendency for individuals with less distinct social preferences (i.e. those that spread their contact moments more evenly over social partners) to reach more patches (Supplementary Table S4). This was particularly the case for females in single-sex compositions, but not for individuals in other sex compositions (Supplementary Table S4).

**Figure 4.**
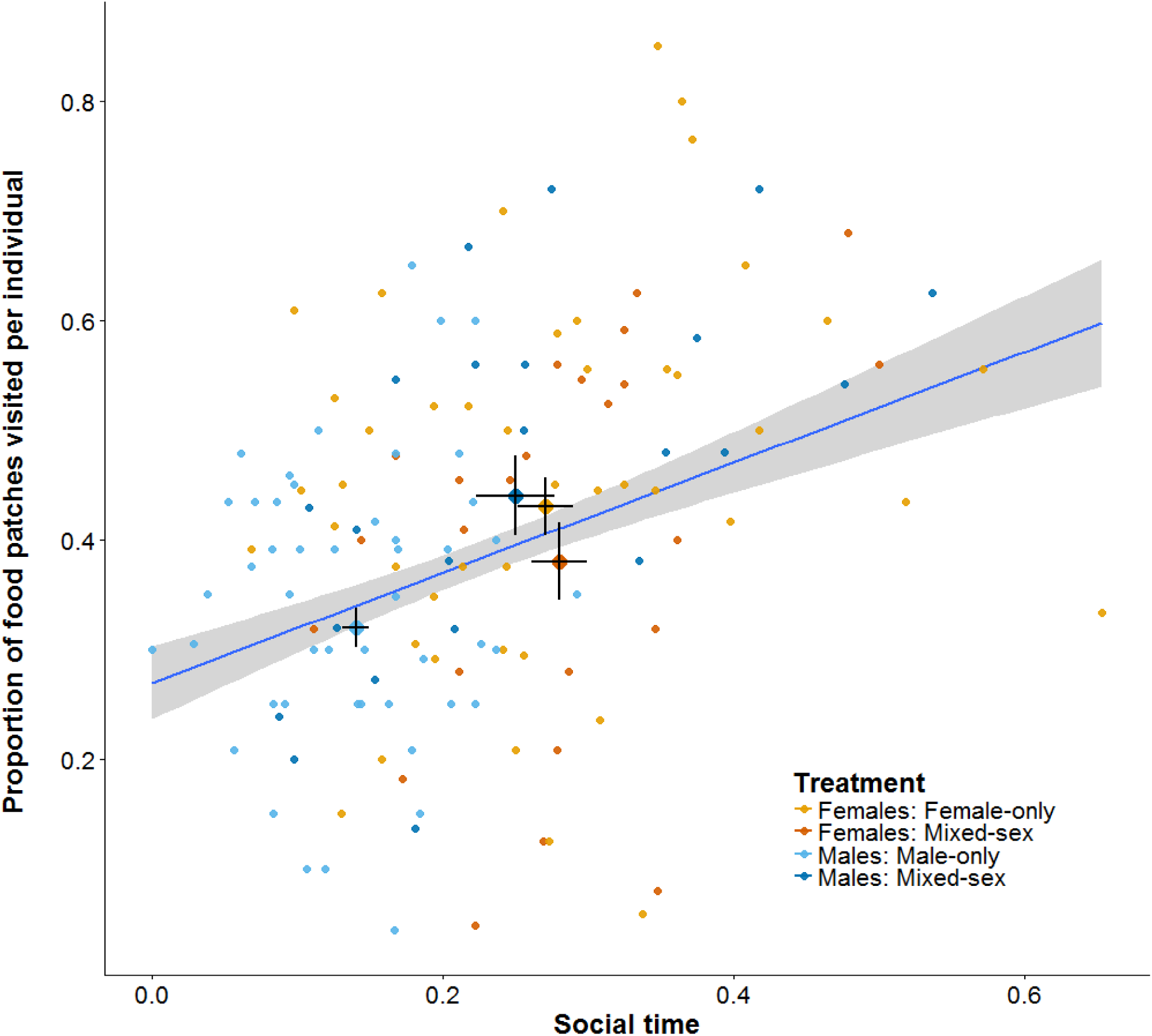
The proportion of novel food patches discovered in relation to individual Social time. A higher Social time value indicates a stronger propensity to spend time in proximity of conspecifics (before the foraging trials). Fish that spend more time social find more novel food patches. Larger points with horizontal and vertical bars indicate observed treatment group means ± 1 SE. Regression lines and 95% CI (shaded area) are based on fitted model values.

### Evidence for sex-biased social attraction

Surprisingly, before the foraging trials, neither males nor females in mixed-sex compositions were more likely than chance to have a female as social partner (Supplementary Table S5). This suggests that males, for which social time was substantially higher in mixed compared to single-sex compositions (Fig. 2), increase their social time with both sexes when females are present. When specifically investigating the social associations within mixed-sex compositions based on the sex of the social partners, male-male associations showed a higher probability of ending (leaving the nearest neighbor) compared to female-female associations (Supplementary Table S6). Male-female (or female-male) association probabilities overlapped with both male and female same-sex probabilities (Supplementary Table S6). A similar trend was visible for the probabilities of individuals to transition from a social state to being alone, but not vice versa (Supplementary Table S6).

During the foraging trials, sex-biases in social attraction were more apparent. In mixed compositions, a smaller proportion of fish joined at a novel food patch if the first individual to arrive was a male compared to a female (*Est ± SE* = −0.60 ± 0.27, *χ*^2^ = 4.95, *N* = 138, *N* individuals = 42, *P* = 0.03). Specifically, a smaller proportion of the males joined (*Est ± SE* = −0.64 ± 0.26, *χ*^2^ = 5.85, *N* = 138, *P* = 0.04), whereas females did not appear to discriminate based on the sex of the first fish at a novel patch (*Est ± SE* = −0.29 ± 0.24, *χ*^2^ = 1.48, *N* = 138, *P* = 0.22). Body size of the first fish did not influence the proportion of batch members that would join (*Est ± SE* = 0.07 ± 0.09, *χ*^2^ = 0.51, *N* = 138, *N* individuals = 42, *P* = 0.47).

## Discussion

Despite great research interest into the evolution and maintenance of social living and foraging, few studies to date have directly manipulated the social environment of foragers in the wild. By effectively manipulating the sex composition, we show that sex composition can be a relevant determinant of individual foraging success in a wild fish population. Male guppies clearly benefitted from foraging with females while female guppies, against our expectations, experienced similar foraging success with or without males. Our study provides unique experimental evidence from the wild on how individual-level traits can interact with population-level traits to shape the adaptive value of social foraging.

Finding novel resources is key to animals foraging in dynamic environments, such as tropical rainforests. Individuals can improve the localization of new resources via specialized behaviours, such as specific movement or search strategies (Humphries *et al.* 2012) or increased sensitivity to certain environmental cues (Dahmani *et al.* 2018). Such strategies might differ between the sexes (Croft *et al.* 2003a), but are unlikely to explain our findings. Males and females were equally successful in initially detecting the food patches (i.e. being the first), both at a group-level, as well as at an individual-level. Instead, we here provide multiple lines of evidence to support that the effects of sex-composition on individual patch discovery were modulated by changes in social dynamics.

We demonstrated that sex-composition affects how much time individuals spent near conspecifics and revealed the effect to be sex-specific, with males, but not females, varying in their time spent social, depending on the presence or absence of the other sex. There are several, not mutually exclusive mechanisms to explain these findings. Males might have become more social in the presence of females in an effort to solicit mating opportunities with females and to engage in agonistic interactions with other males. Consequently, males might have ended up at food patches primarily motivated by the mere presence of conspecifics (i.e. local enhancement (Reader *et al.* 2003; Webster & Laland 2013)). Alternatively, or additionally, males in single-sex compositions may have experienced higher levels of aggression (Magurran & Seghers 1991), causing them to both spend less time social (i.e. avoidance of dominant males) and reach fewer patches (i.e. competitive exclusion), compared to other sex-compositions.

There are also sex-independent mechanisms via which spending more time social can increase an individual’s chances of finding food, ranging from social facilitation to information sharing (Giraldeau & Caraco 2000). For example, spending time near conspecifics likely results in a greater exposure to social information (Danchin *et al.* 2004; Duboscq *et al.* 2016). Indeed, there is a wide variety of species, including tit species (Family: *Paridae*), humpback whales (*Megaptera novaeangliae*) and California sea lions (*Zalophus californianus*), in which social connections predict how novel (foraging) information ‘flows’ through a population (Aplin *et al.* 2012; Allen *et al.* 2013; Schakner *et al.* 2017). Though it is important to note that such correlations do not necessarily reflect active information transfer, i.e. it could reflect individuals regularly being at the same place at the same time (Atton *et al.* 2012; Hasenjager & Dugatkin 2016). Alternatively, being more social might be linked to increased social responsiveness (Kurvers *et al.* 2014; Wolf & Krause 2014), including responsiveness to social foraging cues (Webster & Laland 2011). Future experimental studies manipulating social foraging cues and monitoring the responses of individuals with various levels of ‘social time’ will be necessary to reveal if social responsiveness (or social ‘awareness’) is an underlying mechanism of increased foraging success.

Against our expectations, females did not show a significant reduction in patch discovery in the company of males. This was surprising, because we expected females to be overall less social in the presence of males (see (Wilson *et al.* 2015), avoiding them to prevent sexual harassment (Croft *et al.* 2006; Darden *et al.* 2009), consequently finding fewer patches. Yet, females in mixed compositions were not less social compared to the other compositions. In a previous study, female guppies actively preferred areas with lower male presence even if these areas were associated with higher predation risk (Darden & Croft 2008). Yet, in our study, females with males in their local environment were not less social than females without males in their vicinity. Moreover, females also did not show a distinct preference for socializing with other females over males nor were they more likely to ‘follow’ a first female over a first male at a food patch. Possibly, foraging females in resource limited habitats, like our study site, are more tolerant towards males, also because males, in such habitats, may spend relatively more time finding food and less time harassing females. Resource limited habitats may thus both enhance the need for social foraging as well as its effectiveness by increasing social tolerance between the sexes. A cross-population (cross-habitat) experiment would be an exciting next step to test this hypothesis.

In conclusion, we experimentally demonstrated how the interaction between sex and sex composition can influence the adaptive benefits of social living. Emphasis of future research should be on examining other individual traits (e.g. personality) that may influence social attraction and responsiveness in interaction with composition and on quantifying the relative adaptive benefits of social foraging under different environmental scenarios, for example by varying levels of resource abundance, predictability and predation risk. Indeed, the Trinidadian guppy study system would allow ecologists to examine and effectively manipulate such key environmental factors and thus to more thoroughly understand the driving forces of foraging success in the wild.

## Supporting information

## Acknowledgements

We are grateful to Sergio García Martín and Chente Ortiz for assistance with the video analysis and to Ines Schön and Rebecca Kettwig for assistance with the data collection. L.S. was funded by an IGB Postdoc Fellowship (2017) and an Alexander von Humboldt Postdoc Fellowship (2018).

## Supplementary material includes

Supplementary methods, Supplementary Tables 1-6 and Supplementary Figure 1.

