## Supplementary material for "Females facilitate male patch discovery in a wild fish population"

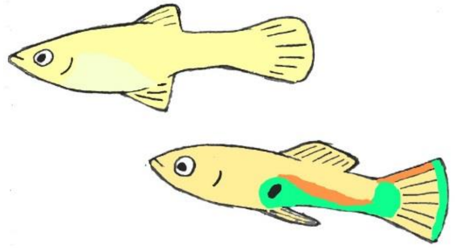

### Supplementary Methods

#### Randomization of social times in the binary model for patch discovery

To examine if variation in individual patch discovery may be explained by corresponding variation in social time, we used the binary model for patch discovery explained in the main text and evaluated the significance of social time via a randomization procedure. More specifically, we permuted the social times (together with the corresponding body lengths) of the individuals. However, while the social times within each batch are highly dependent, the social times of individuals from different batches are not dependent. To avoid mixing the values of dependent and independent variables, we only permuted individual social times within each batch and swapped the individual social times of complete batches, but never swapped individual social times across batches. In order to swap complete batches, all batches need to have the same number of (eight) individuals. Therefore, for 'missing' individuals (5/144) we included dummy values based on the average value for the specific batch.

#### Additional information on the *ϒ*-measure

To quantify the degree to which individuals have social preferences, we calculated the ϒ-measure as the sum of squares of the normalized association strengths (relative number of contact moments) between one individual and all others. The ϒ-measure is sensitive for differences in group size. During our observations of social phenotypes, we had 16 batches of 8 individuals and 2 batches of 7 individuals (Supplementary Table 1b). Therefore, we ran the analyses with ϒ-measure values corrected for group size. We did not change the ϒ-values for the groups with 8 individuals but adjusted the ϒ-values for the individuals in batches with 7 individuals. From the definition of the ϒ-measure it follows that the smallest possible values in a group of 7 ("G7") and in a group of 8 ("G8") differ by a factor of 6/7 = 0.857, more precisely G8 = 0.857 * G7, while the largest possible ϒ-value (= 1) is the same for all group sizes. Therefore, the factor needed for the adjustment lies in the range 0.857 - 1. A previous study on guppies[1] showed that the mean ϒ-values in randomized networks with 7 and 8 individuals in the absence of individual preferences differed by a factor of 0.862, and that a reasonable factor computed from real data was 0.895. We tried out different factors and found that the P-values were almost the same. Therefore, we only reported the results for the factor 0.895.

#### Sex-specific social times

The Markov-chain model introduced by Wilson et al. (2014)[2] consisted of two behavioral (social) states, "being together with another individual" (within four body lengths), denoted by $s$, and "being alone", denoted by $a$. From the transition probabilities between these states, the overall time spent in these states can be concluded by computing the model's stationary distribution. For our analysis of sex-specific attraction in the mixed-sex compositions we split the state $s$ into two states $s_f$ and $s_m$, meaning "being together with a female" and "being together with a male", respectively. By constructing two models, one for female focal fish and one for male focal fish, we computed the overall times spent in these two states depending on the sex of the focal fish. These models were used to estimate the transition probabilities and social times in Tables S5 and S6.

The percentages of the number of contact phases between male-male, female-female, and mixed-sex associations were as one would theoretically expect in the absence of sex-biased social attraction (Supplementary Table S6). This means that the slightly larger individual social time of females in the mixed-sex compositions (28% compared to 26% for males) is mainly caused by the longer duration of female-female contacts. The reciprocal values of the transition probabilities are proportional to the mean durations of contact phases between two individuals, which means that the female-female associations were on average longer than male-male associations (by about a quarter of our 10 s observation interval).

### Supplementary Tables

**Supplementary Table S1a.** Summary of subject sample size. Social observations are missing for one of the females.

| Sex-composition | Females | Males | Total | Notes |
|---|---|---|---|---|
| Single sex | 46 | 48 | 94 | 2 females went missing |
| Mixed sex | 24 | 22 | 46 | 2 males went missing |
| Total | 70 | 70 | 140 | |

**Supplementary Table S1b.** Sample size during social observations. # = Total number of fish present per batch.

| | Pool | | | | | | |
|---|---|---|---|---|---|---|---|
| Sex-composition | 1 | | 2 | | 3 | | Total |
| Female | 8 (9)* | 8 | 7 | 8 | 8 | 7 (8)** | 46 |
| Male | 8 | 8 | 8 | 8 | 8 | 8 | 48 |
| Mixed | 8 | 8 | 8 | 8 | 8 | 7*** | 47 |

**Supplementary Table S1c.** Sample size during foraging trials. # = Total number of fish present per batch.

| | Pool | | | | | | |
|---|---|---|---|---|---|---|---|
| Sex-composition | 1 | | 2 | | 3 | | Total |
| Female | 8* | 8 | 7 | 8 | 8 | 7 (8)** | 46 |
| Male | 8 | 8 | 8 | 8 | 8 | 8 | 48 |
| Mixed | 8 | 8 | 8 | 8 | 8 | 6*** | 46 |

**Supplementary Table S1d.** Sample size of combined social observations and foraging trials. # = Total number of fish present per batch.

| | Pool | | | | | | |
|---|---|---|---|---|---|---|---|
| Sex-composition | 1 | | 2 | | 3 | | Total |
| Female | 7 | 8 | 7 | 8 | 8 | 7 | 45 |
| Male | 8 | 8 | 8 | 8 | 8 | 8 | 48 |
| Mixed | 8 | 8 | 8 | 8 | 8 | 6 | 46 |

* Female present from an earlier batch, not used as social subject but used as foraging subject. Another female that was present during social observations went missing.
** One female turned out to be a male, present both during social network observations and during foraging trials, but not used as subject.
*** One fish disappeared before social observations and another fish went missing before the foraging trials.

**Supplementary Table S2.** Estimated transition probabilities between social states and resulting time spent social for individuals in single-sex or mixed-sex compositions. Non-overlapping confidence intervals indicate significant differences between groups.

| Treatment | Leave Nearest Neighbour | Switch Nearest Neighbour | Social to Alone | Alone to Social | Time social |
|---|---|---|---|---|---|
| | *Mean (95% CI)* | *Mean (95% CI)* | *Mean (95% CI)* | *Mean (95% CI)* | *Percentage* |
| **Female-only** | 0.54 (0.51 - 0.58) | 0.16 (0.14 - 0.18) | 0.38 (0.35 - 0.42) | 0.14 (0.13 - 0.16) | 27 % |
| **Male-only** | 0.83 (0.79 - 0.86) | 0.14 (0.11 - 0.18) | 0.68 (0.64 - 0.73) | 0.11 (0.10 - 0.12) | 14 % |
| **Mixed: Males** | 0.64 (0.59 - 0.68) | 0.16 (0.13 - 0.20) | 0.47 (0.43 - 0.52) | 0.17 (0.15 - 0.19) | 26 % |
| **Mixed: Females** | 0.59 (0.55 - 0.63) | 0.14 (0.11 - 0.17) | 0.45 (0.41 - 0.50) | 0.17 (0.15 - 0.20) | 28 % |

**Supplementary Table S3.** Results of model randomization tests for the relationship between the time spent social and patch discovery. In each of the 10,000 randomization steps we permuted the social times and the corresponding body lengths between individuals within a batch and additionally swapped complete batches. The coefficient of 'social time' (scaled and centered) in the GLMMs (here 'obs. score') was used as test statistic.

| Dataset | *N indiv.* | *# indiv. added* | *% indiv. added* | *obs. score* | *P* |
|---|---|---|---|---|---|
| **All individuals** | 144 | 5 | *3.5%* | 0.220 | **0.0009** |
| **Female-only** | 48 | 3 | *6.3%* | 0.090 | 0.253 |
| **Male-only** | 48 | . | . | -0.039 | 0.636 |
| **Mixed: Females** | 24 | . | . | 0.342 | **0.034** |
| **Mixed: Males** | 24 | 2 | *8.3%* | 0.305 | **0.043** |

**Supplementary Table S4.** Results of model randomization tests for the relationship between spread of social contacts (*ϒ*-measure) and patch discovery. We used the same model as for the analysis of social time, but added the covariate ϒ-measure. In each of the 10,000 randomization steps we permuted the *ϒ*-measures, social times and body lengths between individuals within a batch and additionally swapped complete batches. The coefficient of ϒ-measure (scaled and centered) in the GLMM's (here 'obs.score') was used as test statistic.

| Dataset | *N indiv.* | *# indiv. added* | *% indiv. added* | *obs. score* | *P* |
|---|---|---|---|---|---|
| **All individuals** | 144 | 5 | *3.5%* | -0.085 | 0.080 |
| **Female-only** | 48 | 3 | *6.3%* | -0.363 | **0.001** |
| **Male-only** | 48 | . | . | -0.025 | 0.383 |
| **Mixed: Females** | 24 | . | . | 0.014 | 0.518 |
| **Mixed: Males** | 24 | 2 | *8.3%* | 0.012 | 0.523 |

**Supplementary Table S5.** Observed and random values for the percentages of time spent social with females (F) and males (M), depending on focal sex in the five mixed-sex batches with four males and four females. The randomizations (10,000 repetitions) were performed by randomizing the contact partners of the focal fish within each batch. Non-overlapping confidence intervals indicate significant differences between groups. None of the differences between the observed and random values were significant.

| | **Observed** | | **Randomized** | | **Randomized** | |
|---|---|---|---|---|---|---|
| | Partner F | Partner M | Partner F *(Mean (95% CI))* | | Partner M *(Mean (95% CI))* | |
| Focal F | 13.3% | 15.4% | *12.3%* | *(10.4%, 14.2%)* | *16.4%* | *(14.5%, 18.3%)* |
| Focal M | 16.3% | 10.2% | *15.2%* | *(13.5%, 16.8%)* | *11.4%* | *(9.7%, 13.1%)* |

**Supplementary Table S6.** Estimated probabilities of starting and ending social contact between individuals of the same sex (female-female and male-male, respectively) and of opposite sex (male-female or female-male) for fish in mixed-sex compositions. Additionally, the observed numbers of contact phases and their theoretically expected distribution in the absence of sex-biased social attraction is shown (given that 3/7 of the contacts should be between individuals of same sex and 4/7 between individuals of opposite sex). One group in the mixed sex-treatment was excluded because it only contained three males. Non-overlapping confidence intervals indicate significant differences between groups.

| **Association type** | **Leave Nearest Neighbor** | **Social to Alone** | **Alone to Social** | **Number of contact phases** | |
|---|---|---|---|---|---|
| | *Mean (95% CI)* | *Mean (95% CI)* | *Mean (95% CI)* | *Observed* | *Theoretically expected* |
| **Female-Female** | 0.53 (0.46 - 0.61) | 0.52 (0.44 - 0.59) | 0.08 (0.07 - 0.10) | 118 | 117 |
| **Male-Male** | 0.71 (0.63 - 0.78) | 0.67 (0.59 - 0.74) | 0.08 (0.06 - 0.09) | 117 | 117 |
| **Male-Female / Female-Male** | 0.61 (0.56 - 0.65) | 0.52 (0.47 - 0.56) | 0.10 (0.09 - 0.11) | 310 | 311 |

### Supplementary Figures

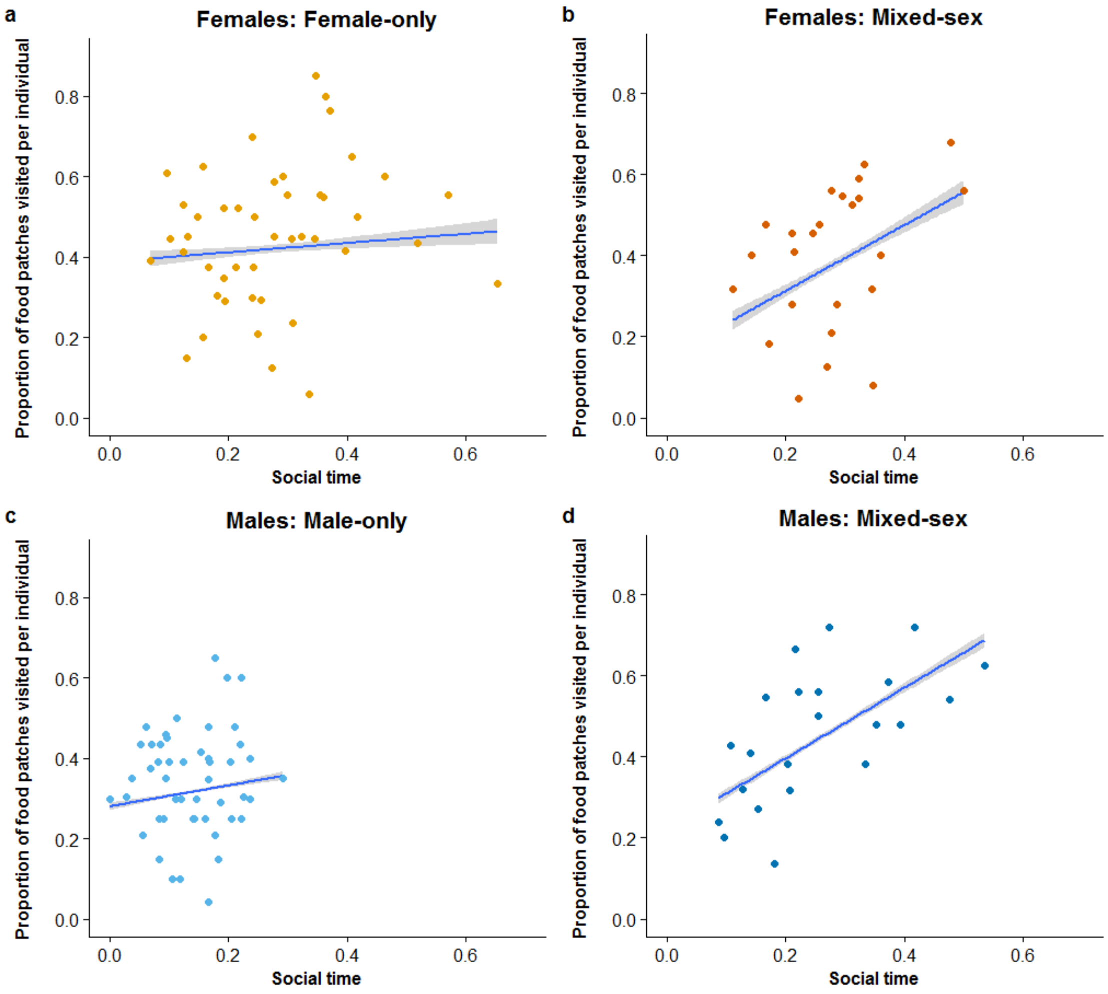


**Supplementary Figure S1. The proportion of food patches discovered by males and females in relation to individual Social time for different sex compositions.** A higher Social time value indicates a stronger propensity to spend time in proximity of conspecifics (before the foraging trials). Males and females that spend more time social in mixed-sex compositions (panel b & d) find more novel food patches, while the correlation for single-sex compositions was non-significant (panel a & c). Regression lines and 95% CI (shaded area) are based on the respective fitted model values.
